# Degradation levels of continuous speech affect neural speech tracking and alpha power differently

**DOI:** 10.1101/615302

**Authors:** Anne Hauswald, Anne Keitel, Ya-Ping Chen, Sebastian Rösch, Nathan Weisz

## Abstract

Understanding degraded speech, e.g. following a hearing damage, can pose a challenge. Previous attempts to quantify speech intelligibility in neural terms have usually focused on one of two measures, namely low-frequency speech-brain synchronization or alpha power modulations. However, reports have been mixed concerning the modulation of these measures, an issue aggravated by the fact that they have normally been studied separately. Using a parametric speech degradation approach, we present MEG studies that overcome this shortcoming. In a first study, participants listened to unimodal auditory speech with three different levels of degradation (original, 7-channel and 3-channel vocoding). Intelligibility declined with declining clarity, implemented by fewer vocoding channels but was still intelligible to some extent even for the lowest clarity level used (3-channel vocoding). Low-frequency (1-7 Hz) speech tracking suggested a u-shaped relationship with strongest effects for the medium degraded speech (7-channel) in bilateral auditory and left frontal regions. To follow up on this finding, we implemented three additional vocoding levels (5-channel, 2- channel, 1-channel) in a second MEG study. Using this wider range of degradation, the speech-brain synchronization showed a similar pattern for the degradation levels used in the first study but further shows that when speech becomes unintelligible, synchronization declines again. The relationship differed for alpha power, which continued to decrease across vocoding levels reaching a floor effect for 5-channel vocoding. Predicting subjective intelligibility based on models either combining both measures or each measure alone, showed superiority of the combined model. Our findings underline that speech tracking and alpha power are modified differently by the degree of degradation of continuous speech but together contribute to the subjective understanding of speech.

## 1. Introduction

Understanding speech can be challenging in normal acoustic environments (e.g. background noise) or due to hearing damage. To compensate for the inferior signal-to-noise ratio of the acoustic information reaching the central auditory system an effortful process is necessary. Indeed, subjective listening effort has been shown to increase with concurrent background noise or competing speakers for speech sounds with fewer acoustic details or lower predictiveness (Wöstmann et al. 2015). “Listening effort” however from a conceptual perspective is not so straightforward, often describing a combination of cognitive demand (usually due to challenging listening situations) and affective-motivational aspects (Peelle 2018). A related compensatory process may be the allocation of increased attentional resources to the incoming (behaviourally relevant) sounds (Wild et al. 2012). Interestingly, both - i.e. the broader concept of listening effort as well as selective attention - have been linked to neural oscillations in the alpha (8-12 Hz) frequency range (e.g. Obleser et al. 2012; Frey et al. 2014; Dimitrijevic et al. 2019). Most studies have reported an alpha power increase in response to degraded speech. This modulation occurs when short degraded stimuli are used (Obleser and Weisz 2012; Obleser et al. 2012). However, in the rare-albeit more naturalistic-situation where sentences have been used, alpha power seems to show the opposite pattern (McMahon et al. 2016; Miles et al. 2017).

Besides the induced neuronal responses broadly linked to the task demands, listening to speech also elicits temporal synchronization of auditory cortical activity to the speech sound. Different frequency bands have been assigned to carry different information with regards to speech signal, with a dominance of delta (1-4 Hz) and theta frequencies (4-7 Hz) capturing phrasal and syllable structure respectively (Greenberg 1998; Poeppel 2003). Synchronization between speech and brain signals is often called neural speech entrainment or speech tracking (see however Alexandrou et al. 2018). Different measures can be used for quantification such as coherence (Hauswald et al. 2018), mutual information (Gross et al. 2013; Keitel et al. 2018), inter-trial correlation (Ding et al. 2014), dissimilarity index (Luo and Poeppel 2007), or temporal response functions (TRF, Ding and Simon 2011; Crosse et al. 2016). Just as alpha power, low frequency speech tracking is modulated by degradation of the speech signal with studies providing mixed findings: Reduced synchronization in this frequency range is linked to reduced intelligibility either operationalized via vocoding (Luo and Poeppel 2007; Ding et al. 2014), time-reversed presentation (Howard and Poeppel 2010; Gross et al. 2013), speech-in-noise (Dimitrijevic et al. 2019), or transcranial electrical stimulation (Riecke et al. 2018; Zoefel et al. 2018). However, using other measures or experimental procedures the opposite pattern has also been shown: e.g. using TRF yields higher M50 and delta synchronization is enhanced for degraded stimuli compared to unaltered stimuli in quiet environment (Ding et al. 2014) and non-native speakers show higher speech entrainment than native-speakers (Song and Iverson 2018). Interestingly the latter observation has also been linked to an increase of listening effort. To complicate things further, multi-speaker and auditory spatial attention studies using sentences or narratives have repeatedly found stronger speech tracking (delta and theta band) for attended compared to unattended speech (Ding and Simon 2012; Rimmele et al. 2015; Viswanathan et al. 2019) in auditory cortices and areas in the vicinity thereof (Horton et al. 2013; Zion Golumbic et al. 2013).

Thus both relevant measures -i.e. speech tracking and alpha power- are frequently however not consistently linked with listening effort (Song and Iverson 2018; Dimitrijevic et al. 2019), selective attention (Frey et al. 2014; Rimmele et al. 2015) or intelligibility of the stimuli (Vanthornhout et al. 2019). A problem contributing to this situation is that very few studies have investigated these measures simultaneously. One study, using a speech-in-noise task, reported decreasing speech tracking and increasing alpha power in response to increasing listening effort in cochlear implant users (Dimitrijevic et al. 2019). However, here again, short stimuli (digits) were presented which is a paradigm rather remote from real-life listening situations. We present findings from two MEG studies that together overcome this situation using continuous speech and a wide range of degradation levels. Importantly derived from the same data set, we show a differential modulation pattern of both measures: Speech tracking increases the stronger stimuli are degraded as long as some intelligibility is still warranted, to then decrease beyond this critical point. Alpha power on the other hand decreases with increased degradation and stays low even when unintelligible. Using linear mixed effects models, we show that combining speech tracking and alpha power is superior in predicting subjective intelligibility of degraded speech, as compared to models based on one of the neural measures alone.

## 2. Study 1

### 2.1 Material and Methods

#### 2.1.1 Participants

Twenty-eight individuals participated in the study (female=17, male=11). Mean age was 23.82 years (*SD* = 3.712), with a range between 19 and 37 years. We recruited only German native speakers and people who were eligible for MEG recordings, i.e. without non-removable ferromagnetic metals in or close to the body. Participants provided informed consent and were compensated monetarily or via course credit. Participation was voluntary and in line with the declaration of Helsinki and the statutes of the University of Salzburg. The study was preregistered at OSF (https://osf.io/dpt34/).

#### 2.1.2 Stimuli

For the MEG recording 12 audio files were created from audio-visual recordings of a female speaker reading Goethe’s “Das Märchen” (1795). Stimuli lengths varied between approximately 30 s and 3 min, with two stimuli of 15 s, 30 s, 60 s, 90 s, 120 s, 150 s, and 12 of 180 s. Each stimulus ended with a two-syllable noun within the last four words. In order to keep participants’ attention on the stimulation, we asked participants after each stimulus to choose from two presented two-syllable nouns the one that had occurred within the last four words of a sentence. The syllable rate of the stimuli varied between 4.1 and 4.5 Hz with a mean of 4.3 Hz.

Noise-vocoding of all audio stimuli was done using the vocoder toolbox for Matlab (Gaudrain, E. 2016) and we created conditions with 7 and 3 channels (fig. 1A). For the vocoding, the waveform of each audio stimulus was passed through two Butterworth analysis filters (for 7 and 3 channels) with a range of 200-7000 Hz, representing equal distances along the basilar membrane. Amplitude envelope extraction was done with half-wave rectification and low-pass filtering at 250 Hz. The envelopes were then normalized in each channel and multiplied with the carrier. Then, they were filtered in the band, and the RMS of the resulting signal was adjusted to that of the original signal filtered in that same band. Auditory stimuli were presented binaurally using MEG--compatible pneumatic in--ear headphones (SOUNDPixx, VPixx technologies, Canada). The trigger-sound delay of 16 ms was measured (The Black Box Toolkit v2) and corrected for during preprocessing.

**Figure 1:**
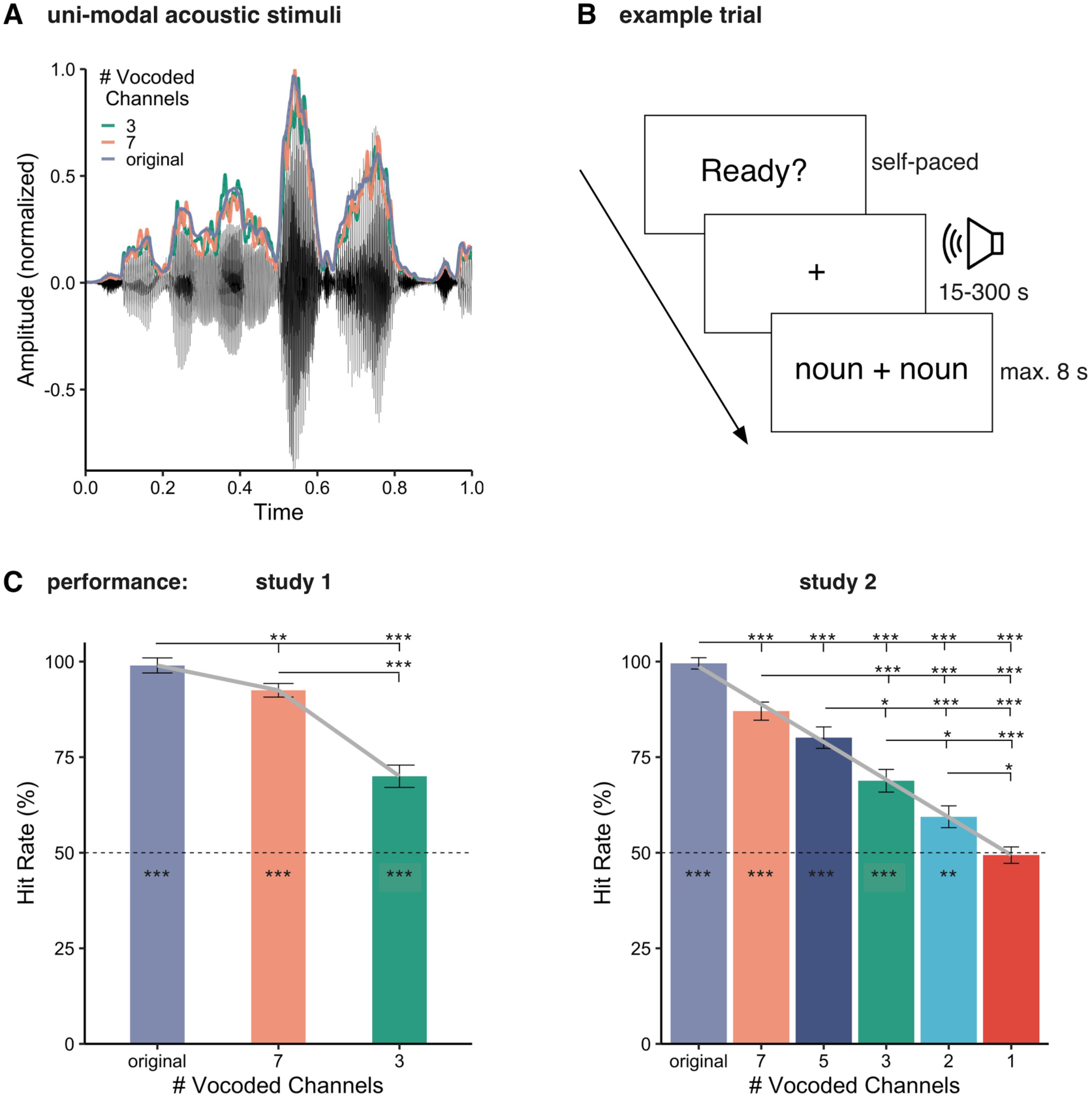
**A**) an exemplary audio file with the corresponding envelope as well as with the envelopes of the vocoded audio stimuli presenting either 7 or 3 channels as used in study 1. **B**) Example trial of unimodel acoustic stimulation. Participants started the presentation self-paced and listened to the stimulus during the visual presentation of a fixation cross. When the stimulus ended, participants were presented with two nouns of which they had to pick the one they perceived in the sentence before. **C**) hit rates in the behavioral experiment in study 1 and 2 using acoustic stimuli of single sentences (range of 2-15 s). The grey curves represent the model-based predicted behavioral response (left: model combining linear and quadratic term; right: linear model). Bars represent standard error, *p_fdr*<0.05*, *p_fdr*<0.01**, *p_fdr*<0.001***.

In the experiment, in addition to the unimodal auditory stimuli also uni-modal visual stimuli were presented, which will not be reported or discussed here as the visual degradation manipulation was not comparable to the acoustic one. The unimodal stimuli were presented to the participants in three consecutive audio-only blocks and three consecutive video-only blocks via in-ear-phones and a projector system respectively. The order of video and audio blocks was balanced. Each block contained 4 stimuli which were presented either in an unaltered version or in one of the two degraded versions. The order of the stimuli was random and did not follow the order of the original story. The assignment of stimuli to conditions was controlled in order to obtain similar overall length of stimulus presentation (approx. 400 s) for each modality and degradation levels. We instructed participants to attend to the speech which they would either see or hear. In order to keep participants’ attention on the stimulation a behavioral response was required after each stimulus. At the end of each stimulus, a target and a distractor word would appear next to each other. The participants were asked to decide which of the words was presented as the last noun and within the last four words by pressing the button on the side of the response pad that matched the presentation side of the word they chose (fig. 1B). Presentation side of target and distractor words was random. Following the response, they could self-initiate the next trial via a button press. Each block was followed by a short self-determined break. This procedure resulted in only four responses per condition and therefore we added a behavioral experiment following all six blocks, to assess performance. Responses were acquired via a response pad.

For this additional behavioural experiment, we used a total of 24 unimodal audio stimuli of a different female speaker reading Antoiné St. Exupery’s “The little prince” (1943). Each stimulus contained a single sentence (length between 2 and 15 seconds) with a two-syllable noun (target word) within the last four words. We created a list of different two-syllable nouns (distractor words) which we also drew from “The little prince” but were not presented during the stimulation. Similar to the main experiment, participants had to choose between two alternatives and the chance level was 50%. The behavioral stimuli were manipulated in the same way as the stimuli for the MEG experiment. Stimulus presentation was controlled using in-house wrapper (https://gitlab.com/thht/th_ptb) for the MATLAB-based Psychtoolbox (Brainard 1997; Pelli 1997; Kleiner et al. 2007).

#### 2.1.3 Data acquisition and analyses

##### 2.1.3.1 Extraction of acoustic speech envelope

We extracted the acoustic speech envelope using the Chimera toolbox by Delguette and colleagues (http://research.meei.harvard.edu/chimera/More.html) where nine frequency bands in the range of 100 to 10000 Hz were constructed as equidistant on the cochlear map (Smith et al. 2002; Chandrasekaran et al. 2009; Gross et al. 2013). Sound stimuli were band-pass filtered (forward and reverse) in these bands using a 4^th^-order Butterworth filter. For each band, envelopes were calculated as absolute values of the Hilbert transform and were averaged across bands to obtain the full-band envelope that was used for coherence analysis. We did this for all three conditions (original, 7-chan, 3-chan) resulting in virtually identical envelopes for those conditions (fig. 1A).

##### 2.1.3.2 MEG acquisition and preprocessing

Data acquisition and analyses closely resembles with minor exceptions the one described in Hauswald et al. (2018). MEG was recorded at a sampling rate of 1 kHz using a 306-channel (204 first order planar gradiometers) Triux MEG system (Elekta-Neuromag Ltd., Helsinki, Finland) in a magnetically shielded room (AK3B, Vakuumschmelze, Hanau, Germany). The MEG signal was online high-pass and low-pass filtered at 0.1 Hz and 330 Hz, respectively. Prior to the experiment, individual head shapes were digitized for each participant including fiducials (nasion, pre-auricular points) and around 300 points on the scalp using a Polhemus Fastrak digitizer (Polhemus, Vermont, USA). We use a signal space separation algorithm provided by the MEG manufacturer and implemented in the Maxfilter program (version 2.2.15) to remove external noise from the MEG signal (mainly 16.6 Hz, and 50Hz plus harmonics) and realign data to a common standard head position (across different blocks based on the measured head position at the beginning of each block).

Data were analyzed offline using the Fieldtrip toolbox (Oostenveld et al. 2011). First, a high-pass filter at 1 Hz (6th order Butterworth IIR) was applied to continuous MEG data. Then, trials were defined according to the duration of each stimulus and cut into segments of two seconds to increase signal-to-noise ratio. As we were interested in frequency bands below 20 Hz and in order to save computational power, we resampled the data to 150 Hz. Independent component analysis was applied separately for visual and auditory blocks and we then identified components corresponding to blinks and eye movements and cardiac activity and removed them. On average 3.25 (SD: 1.143) components were removed for auditory blocks. Sensor space data were projected to source space using linearly constrained minimum variance beamformer filters (Van Veen et al. 1997) and further analysis was performed on the obtained time-series of each brain voxel (http://www.fieldtriptoolbox.org/tutorial/shared/virtual_sensors in FieldTrip). To transform the data into source space, we used a template structural magnetic resonance image (MRI) from Montreal Neurological Institute (MNI) and warped it to the subject’s head shape (Polhemus points) to optimally match the individual fiducials and headshape landmarks. This procedure is part of the standard SPM (http://www.fil.ion.ucl.ac.uk/spm/) procedure of canonical brain localization (Mattout et al. 2007).

A 3D grid covering the entire brain volume (resolution of 1 cm) was created based on the standard MNI template MRI. The MNI space equidistantly placed grid was then morphed to individual headspace. Finally, we used a mask to keep only the voxels corresponding to the grey matter (1457 voxels). Using a grid derived from the MNI template allowed us to average and compute statistics as each grid point in the warped grid belongs to the same brain region across participants, despite different head coordinates. The aligned brain volumes were further used to create single-sphere head models and lead field matrices (Nolte 2003). The average covariance matrix, the head model and the leadfield matrix were used to calculate beamformer filters (regularization factor of 10%). The filters were subsequently multiplied with the sensor space trials resulting in single trial time-series in source space. The number of epochs across conditions was equalized.

We applied a frequency analysis to data of all three conditions (acoustic: original, 7- chan, 3-chan) calculating multitaper frequency transformation (dpss taper: 1-25 Hz in 1 Hz steps, 3 Hz smoothing). These values were used for the analyses of alpha as well as for the coherence calculation between each virtual sensor and the acoustic speech envelope. For all three conditions we used the envelopes of the original, nonvocoded acoustic signal. Then, the coherence between activity at each virtual sensor and the acoustic speech envelope during acoustic stimulation in the frequency spectrum was calculated and averaged across trials. We refer to the coherence between acoustic speech envelope and brain activity as speech tracking. As a sanity check, we calculated grand averages of the speech tracking of the three conditions to see if they show the expected peak around 4 Hz (fig. 2A).

**Figure 2:**
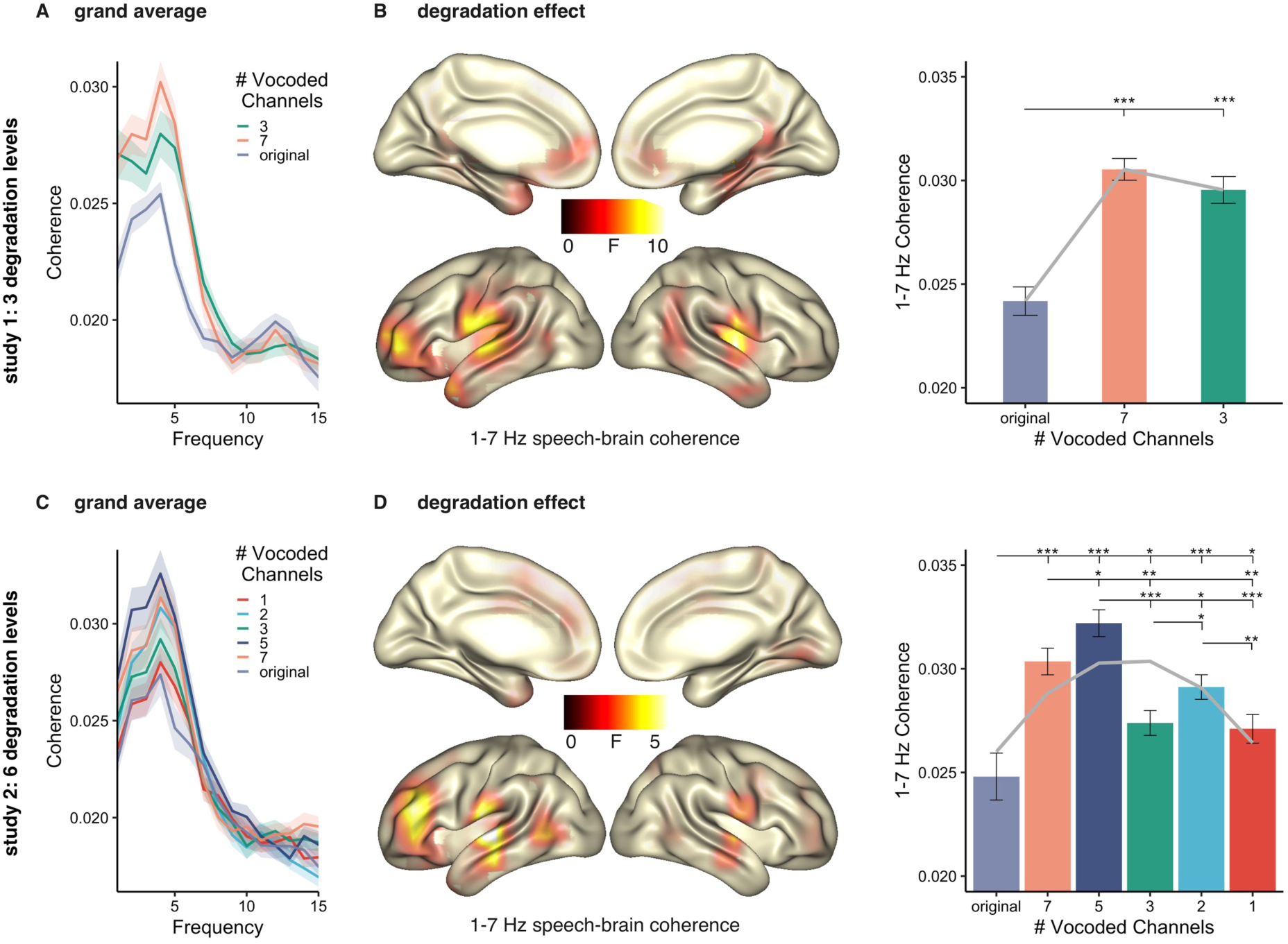
**A**) Frequency spectrum of the speech tracking (coherence) for the three conditions averaged across all voxels. **B) Left**: source localizations of degradation effects on speech tracking (1-7 Hz) during acoustic stimulation across three conditions (original, 7-chan, 3- chan) in bilateral temporal and left frontal regions. **Right**: individual speech tracking values of the three conditions extracted at voxels showing a significant effect contrasted with each other. The grey curve represents the predicted tracking values by the model combining linear and quadratic terms. **C**) Frequency spectrum of the speech tracking for the six conditions averaged across all voxels. **D) Left:** source localizations of degradation effects on speech tracking (1-7 Hz) during acoustic stimulation across six conditions (original, 7- chan, 5-chan, 3-chan, 2-chan, 1-chan) in bilateral temporal and left frontal regions. **Right**: individual speech tracking values of the six conditions extracted at voxels showing a significant effect contrasted with each other. The grey curve represents the predicted tracking values by the model that combines linear and quadratic terms. Bars represent standard error, *p*_*fdr*_<0.05*, *p*_*fdr*_<0.01**, *p*_*fdr*_<0.001***

#### 2.1.4 Statistical analyses

We analyzed the responses from the behavioral experiment. Due to technical problems, behavioral measures are missing for 3 participants and the responses of the remaining 25 participants were analyzed. We used repeated-measures ANOVA to compare across the conditions and then dependent samples t-tests to compare hit rates between conditions and against chance level (50 %).

Most studies on speech-brain entrainment report findings of frequencies below 7 Hz; therefore, we analyzed frequencies between 1 and 7 Hz. For alpha power we analyze 8-12 Hz. For both MEG alpha power and 1-7 Hz coherence data we applied repeated-measures ANOVA (ft_statfun_depsamplesFunivariate in FieldTrip) to test modulations of neural measures across the different degradation levels. To control for multiple comparisons, a non-parametric Monte-Carlo randomization test was undertaken (Maris and Oostenveld 2007). The test statistic was repeated 5000 times on data shuffled across conditions and the largest statistical value of a cluster coherent in source space was kept in memory. The observed clusters were compared against the distribution obtained from the randomization procedure and were considered significant when their probability was below 5%. Effects were identified in source space and the corresponding individual coherence and power values were extracted and post-hoc *t*-tests between conditions were corrected for multiple comparisons by using the FDR method (Benjamini and Hochberg 1995). For visualization, source localizations were mapped onto inflated surfaces as implemented in FieldTrip.

We used linear mixed models to further test how our data (i.e. behavioral response, speech tracking, and alpha power) are influenced by the vocoding levels. At the outset we tested a simple linear model [*recorded measure = (vocoding levels)*] and compared it to a more complex (combined) by adding a quadratic term [*recorded measure = (vocoding levels + (vocoding levels)^2)]*. These two models were compared using an ANOVA test. The respective best model was subsequently reapplied to the data for each individual and the average for these predicted model outcomes are displayed alongside the actual (grand) average results in the relevant bar graphs (grey curves).

### 2.2 Results

#### 2.2.1 Behavioral results

The mean hit rate for original stimuli was 99% (SD: 3.43%) for the original sound files, 92.5% (SD: 10.21%) for 7-chan vocoded stimuli, and 69.44% (SD: 18.75%) for 3-chan vocoded stimuli. A one-way ANOVA across the three conditions revealed a main effect (F(72)=37.14, p=8.28e-12).Comparing the different vocoding levels with each other, showed higher hit rates for nonvocoded stimuli than for 7-chan (t(24)=3.376, *p*_*fdr*_=0.0025) or 3-chan vocoded stimuli (t(24)=7.632, *p*_*fdr*_=1.437e-7). 7-chan had higher hit rates than 3-chan vocoded stimuli (t(24)=6.2354, *p*_*fdr*_=2.8733e-6). All conditions also showed significant above-chance (50%) hit rates (fig. 1C): for nonvocoded stimuli t(24) = 70.787, *p*_*fdr*_ = 1.3341e-28, for 7-chan t(24)=20.821, *p*_*fdr*_=2.1531e-16, and for 3-chan vocoded t(24)=5.333, *p*_*fdr*_=2.1494e-5. The linear mixed models revealed significant linear decrease across conditions (χ^2^ =72.003, p < 2.2e- 16). Adding a quadratic term to the model benefitted the data prediction (*model comparison*: χ^2^ =7.8982, p < 0.004949; grey curve in Fig. 1 C left).

#### 2.2.2 MEG data

##### Degradation related effects

To investigate the effects of reducing the acoustic information, we ran a cluster-corrected repeated measures ANOVA for the speech tracking (1-7 Hz coherence, see spectral distribution in Fig. 2A) of the 3 conditions (original, 7-chan, 3-chan). An effect of degradation between 1-7 Hz (*p=*0.0009) was yielded with maxima in bilateral middle temporal and left frontal regions (Fig. 2B, left). In these areas, the original audio stimuli lead to the weakest speech tracking, while the stimuli with the medium degradation (7-chan) elicited the strongest speech tracking (Fig. 2B, right). Listening to the original audio files elicited lower tracking than listening to the 7-chan (t(27)=-7.798, *p*_*fdr*_=6.58e-8) or 3-chan version (t(27)=-5.593, *p*_*fdr*_=9.33e-6). The two vocoded stimuli classes did not differ significantly (t(27)=1.139, *p*_*fdr*_=0.264). The linear mixed models revealed a significant linear pattern across conditions (χ^2^ =26.868, p=2.179e-07). Adding a quadratic term to the model benefitted the data prediction (*model comparison*: χ^2^=19.998, p=7.751e-06; grey curve in Fig. 2B right).

The same statistical analysis applied to alpha power (8-12 Hz, spectral distribution in Fig. 3A) over original, 7-chan and 3-chan revealed an effect of degradation (*p=*0.0009, fig. 3B), with alpha power during unaltered stimuli being higher during than 7-chan vocoding (t(27)=3.095, *p*_*fdr*_=0.0045) and 3-chan vocoding (t(27)=4.09, *p*_*fdr*_=0.001). Compared to 7-chan vocoding alpha power during 3-chan vocoding decreased even further (t(27)=3.738, *p*_*fdr*_=0.0013). The effect was widespread and covered most of the brain (present in 1357 of 1457 voxel) with a clear maximum in the left angular/parietal inferior cortex. The linear mixed models revealed a significant linear pattern across conditions (χ^2^ =30.292, p=3.716e-08). Adding a quadratic term to the model did not benefit the data prediction (*model comparison*: χ^2^=0.2185, p= 0.6402; grey curve Fig. 3B right).

**Figure 3:**
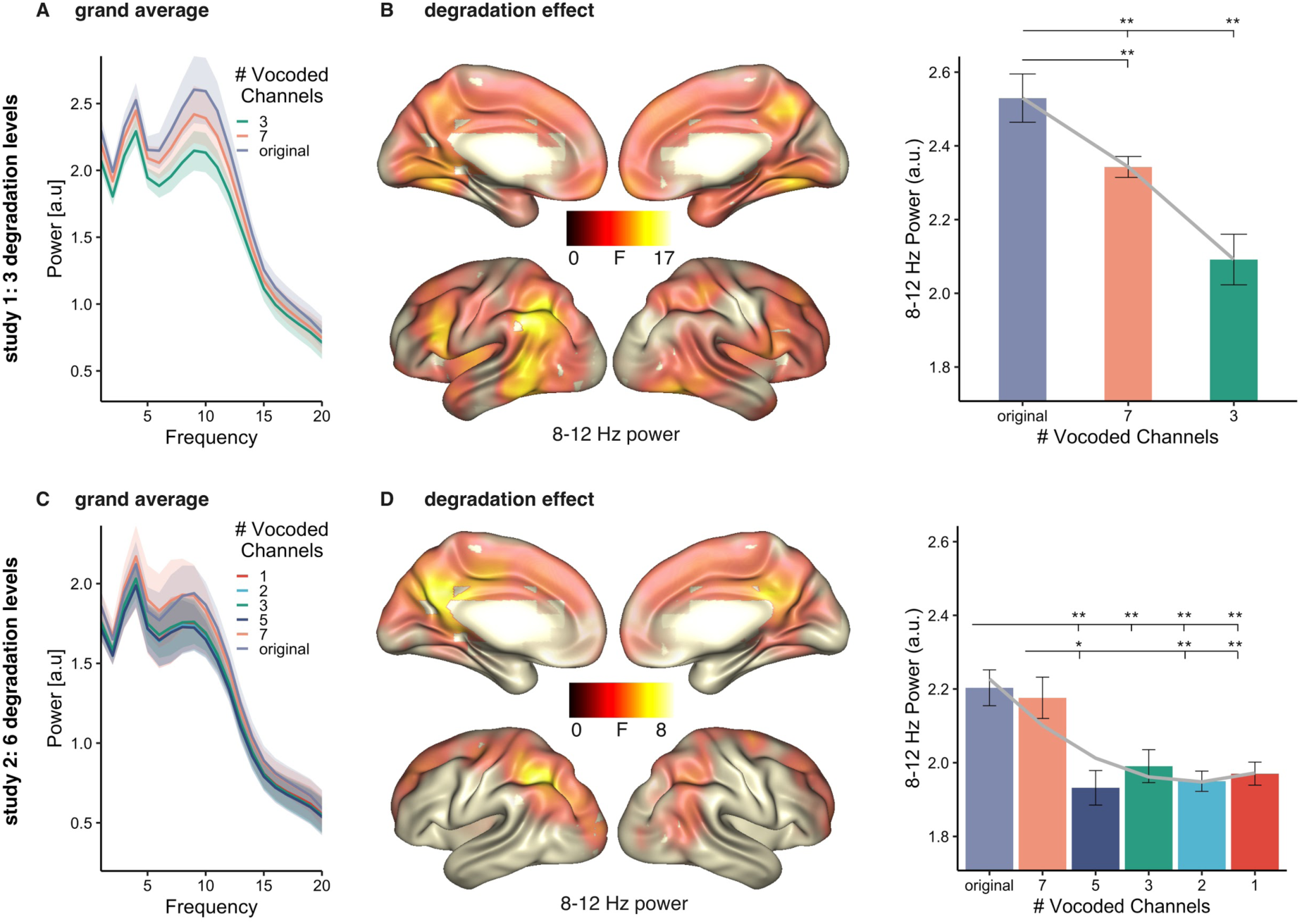
**A**) Frequency spectrum of the power for the three conditions averaged across all voxels. **B) Left**: source localizations of degradation effects on alpha power (8-12 Hz) across three conditions (original, 7-chan, 3-chan) with maxima in left angular gyrus and inferior parietal lobe, left frontal and inferior temporal regions. **Right**: individual 8-12 Hz power values of the three conditions extracted at voxels showing a significant effect contrasted with each other. The grey curve represents the predicted tracking values by the linear model. **C**) Frequency spectrum of the power for the six conditions averaged across all voxels. **D) Left:** source localizations of degradation effects on alpha power (8-12 Hz) across six conditions (original, 7-chan, 5-chan, 3-chan, 2-chan, 1-chan) with maxima in left angular gyrus and inferior parietal lobe. **Right**: individual 8-12 Hz power values of the three conditions extracted at voxels showing a significant effect contrasted with each other. The grey curve represents the predicted tracking values by the model that combines linear and quadratic terms. Bars represent standard error, *p*_*fdr*_<0.05*, *p*_*fdr*_<0.01**, *p*_*fdr*_<0.01***

## 3. Study 2

The findings from study 1 offer two important insights: First, the increase in speech-brain coherence and the decrease in alpha power with decline in acoustic detail is at odds with several previous studies (e.g. Obleser et al. 2012; Dimitrijevic et al. 2019). However, those studies have usually employed very brief stimuli which is uncommon in natural listening situations. Second, the findings suggest that the relationship between degradation and speech tracking might not be linear and possibly behave differently than the relationship between degradation and alpha power. Therefore, we conducted a second study, to replicate the first by using again the previous vocoding levels and further extend it by adding three more vocoding levels: First, we added 5-channel vocoding to fill the gap between 7- and 3-channel vocoding, where comprehension is challenging but still possible. Furthermore, we also added 2- and 1-channel vocoding to make sure we also present unintelligible material.

### 3.1 Materials and Methods

#### 3.1.1 Participants

Twenty-four individuals participated in the second study (female=11, male=13). Mean age was 26.37 years (*SD* = 5.648), with a range between 18 and 45 years. We recruited only German native speakers and people who were eligible for MEG recordings, i.e. without non-removable ferromagnetic metals in or close to the body. Participants provided informed consent and were compensated monetarily or via course credit. Participation was voluntary and in line with the declaration of Helsinki and the statutes of the University of Salzburg.

#### 3.1.2 Stimuli

We used the same auditory stimulus material and experimental design as in study 1, but expanded the degradation levels to include additionally 5-channel, 2-channel and 1- channel vocoding. Overall we had six levels of degradation: original, 7-channel, 5-channel, 3- channel, 2-channel, 1-channel.

#### 3.1.3 Data acquisition and analyses and statistical analyses

All steps of data acquisition, analysis and statistics were identical to study 1. The only exception was that only 17 individuals participated also in the behavioral experiment. An additional 10 participants provided only behavioral data, resulting in 27 data sets of the behavioral part.

### 3.2 Results

#### 3.2.1 Behavioral results

The mean hit rate was 99.46% (SD: 2.61%) for the original sound files, 86.415% (SD: 13.54%) for 7-chan vocoded stimuli, 78.8% (SD: 17.45) for 5-chan, 67.39% (SD: 17.57%) for 3-chan, 56.52% (SD: 15.01) for 2-chan, and 50% (SD: 13.06%) for 1-chan vocoded stimuli. A one-way ANOVA across the three conditions revealed a main effect (F(156)=47.83, p=8.28e- 30). Comparing the different vocoding levels with each other, showed higher hit rates for nonvocoded stimuli than any of the other conditions (all t values > 4.83, all *p*_*fdr*_ values *<* 0.000051). 7-chan vocoded had higher hit rates than 3-, 2-, and 1-chan vocoded stimuli (all t values > 4.96, all *p*_*fdr*_ values *<* 0.000055). 5-chan vocoding had higher hit rates than 3-, 2-, and 1-chan vocoded stimuli (all t values > 2.62, all *p*_*fdr*_ values *<* 0.05). 3-chan vocoding had higher hit rates than 2-, and 1-chan vocoded stimuli (all t values > 2.24, all *p*_*fdr*_ *values <* 0.05). 2-chan had higher hit rates than 1-chan vocoded stimuli (t(26)= 2.74, *p*_*fdr*_ *=* 0.013). The nonvocoded stimuli as well as the 7-chan, 5-chan, 3-chan, and 2-chan vocoded conditions showed significant above-chance (50%) hit rates (all t values > 3.1, all *p*_*fdr*_ *values <* 0.01, fig. 1C, right). The contrast with 1-chan vocoded stimuli did not show a difference (t(26)=-0.25 (*p*_*fdr*_=0.801). The linear mixed models revealed significant linear decrease across conditions (χ^2^ =282.09, p < 2.2e-16). Adding a quadratic term to the model did not result in better prediction of the data (*model comparison*: χ^2^=0.012, p=0.9126; grey curve in Fig. 1 C right).

#### 3.2.2 MEG data

##### Degradation related effects

To investigate the effects of reducing the acoustic information, we ran a cluster-corrected repeated-measures ANOVA for the speech tracking (1-7 Hz coherence, see spectral distribution in Fig. 2C) of the 6 conditions (original, 7-chan,5-chan, 3- chan, 2-chan, 1-chan). An effect of degradation between 1-7 Hz (*p=*0.0009) was located in virtually identical regions as in study 1 (bilateral middle temporal and left frontal regions). In these areas, the original audio stimuli and the most strongly degraded (1-chan) led to the weakest speech tracking, while the stimuli with 5-chan degradation elicited strongest speech tracking (Fig. 2D). Listening to the 5-chan vocoded audio files elicited higher tracking than listening to any of the other conditions (all t values > 2.5, all *p*_*fdr*_ values < 0.05). Listening to the original nonvocoded audio files elicited lower tracking than listening to any of the other conditions (all t values > −2.21, all *p*_*fdr*_ values < 0.05). Similarly, listening to 1-chan vocoded audio elicited lower trakcing than listening to the 7-chan, 5-chan, and 2-chan versions (all t values > −3.88, all *p*_*fdr*_ values < 0.01). Further, 3-chan vocoding yielded lower tracking than 7- chan (t(23)=-3.64, *p*_*fdr*_=0.0026) and 2-chan version (t(23)=-2.72, *p*_*fdr*_=0.02). The linear mixed models did not reveal a linear pattern (χ^2^ =0.1588, p=0.6903). Adding a quadratic term to the model significantly benefited data prediction (*model* comparison: χ^2^=23.642, p=1.16e-06; grey curve in Fig. 2D right).

Calculating cluster-corrected repeated-measures ANOVA for alpha power (8-12 Hz, see spectral distribution in Fig. 3C) over the six conditions revealed an effect of degradation (*p=*0.0009, fig. 3D) with maxima analogous to Study 1, i.e. in left angular gyrus and inferior parietal lobe. Nonvocoded and 7-chan vocoded stimuli eliciting higher alpha power any of the other conditions (all t values > 2.904, all *p*_*fdr*_ values < 0.05) except 7-chan and 3-chan did not show a conclusive difference (t(23)=2.243, *p*_*fdr*_=0.0653). The linear mixed models did reveal a significant linear pattern (χ^2^ =22.206, p=2.449e-06). Adding a quadratic term to the model significantly benefited data prediction (*model comparison*: χ^2^=6.6019, p=0.01019, grey curve in Fig. 3D right).

#### 3.2.3 Using neural measures to predict speech intelligibility

Our MEG data, especially using the richer set of degradation levels in study 2, indicate a differential impact on our neural measures. This should serve as a precaution against simplistically equating the neural measures to such abstract concepts as listening effort. In order to be functionally relevant one would expect that these neural measures predict speech intelligibility. However, based on the previous analysis this is not clear. In a last hypothesis generating step of this study, with the aim of guiding future research, we postulate *alpha* to be an “activation” proxy of neural ensembles. However, such an “activation” may not necessarily lead to activation of veridical (i.e. ∼intelligible) representations (Griffiths et al. 2019) especially when the sound becomes increasingly degraded. We speculate that neural tracking may reflect the outcome of this combination between “activation” and “veridicability”. Since no continuous time-varying quantification of the latter concept is available, behaviorally assessed “intelligibility” can serve as a proxy. The basic assumption of this *combined model* can thus be expressed as:

*(1) Neural Tracking = Activation x Intelligibility*

Thus by reordering (1) we obtain a simple model to predict intelligibility of speech from neural data:

*(2) Intelligibility = Neural Tracking / Activation*

where is the estimated slope. The parameters of the model can be estimated using a linear mixed model (using lme4 library implemented in R, Bates et al. 2015) and the model can be compared to competing models (see below). Models were fit using random intercept and slopes. We used the speech-brain coherence and the alpha power of all significant voxels during the nonvocoded “effortless” condition to normalize the other five challenging (i.e. vocoded) listening conditions. For each participant (17 participants who contributed MEG and behavioral data) we then used the model to estimate *intelligibility* values for the vocoded conditions. This *predicted intelligibility* was then compared to the *observed intelligibility* (behavioral response) (χ^2^ =8.3457, p=0.003866).

In order to evaluate whether a combination between neural tracking and activation yields a benefit we compared predicted intelligibility to two simpler models either using only speech tracking (*tracking model*)

(3) *Intelligibility = Neural Tracking; (*χ^2^ =4.6476, p=0.0311)

or only activation (*activation model*).

(4) *Intelligibility = Activation; (*χ^2^ =2.7638, p=0.09642)

Directly comparing the *combined model* to the *tracking model* and in a separate step to the *activation model* shows superiority of the *combined model* (*combined model* vs *tracking model*: χ^2^ =3.436, p < 2.2e-16; *combined model* vs *activation model*: χ^2^ =5.2411, p < 2.2e-16). This means that speech tracking and alpha power together can better predict the behavioral response than either of the factors alone.

## 4. Discussion

As shown in previous studies (e.g. Luo and Poeppel 2007; Obleser and Weisz 2012; Obleser et al. 2012), listening to degraded speech modulates speech tracking and alpha power. The pattern of this modulation varies across studies suggesting that it might depend on experimental implementation and the two measures are not commonly reported together in the field of degraded speech. We advance these previous findings by investigating the effects of degraded speech stimuli on speech tracking and on alpha power in two studies using continuous speech and various degradation levels. In the first study we used three levels of vocoding. Based on the behavioral results and the MEG findings we conducted a second study expanding the degradation levels with one additional intermediate vocoding level (5-channel) and two very low vocoding levels (1- and 2-channel). As both studies yield very similar results in terms of behavior, speech tracking, and alpha power, we will discuss them together.

### 4.1 Behaviorally assessed intelligibility

To be sure that our manipulation actually affects intelligibility, participants performed a behavioral experiment after the MEG experiments. These were in both cases similar to the MEG experiment (with identical degradation levels) but with shorter stimuli, enabling us to assess more trials. The stimuli varied between 2 and 10 seconds instead of 15 seconds and 3 min as during the MEG recording. The data showed that participants decline in performance when the stimuli are degraded, which is in line with other studies showing a linear decline in performance (Strelnikov et al. 2011; McGettigan et al. 2012). The exact number of channels needed for high performance understanding depends on the stimulus material and the specific experimental setup (Dorman et al. 1997; Loizou et al. 1999). For our first study, we conclude that even the 3-chan condition was challenging yet not completely unintelligible given that performance is still higher than expected by chance. Therefore, we added the two lower vocoding conditions (2-chan, 1-chan) in the second study. Results of study 2 showed again that performance declines with degradation and that complete unintelligibility is reached with 1-channel vocoding.

### 4.2 Speech-tracking across degradation level follows an inverted u-shape

To elucidate if the intelligibility, measured by degradation level, affects the speech tracking, measured by speech-brain coherence, we calculated a repeated-measures ANOVA of the low frequency speech-brain coherence (1-7 Hz) across the three (study 1), respectively six (study 2) conditions. For both studies, this revealed bilateral sources in temporal-including auditory-cortex and left frontal regions in which higher tracking was associated with a medium level of degradation. The quadratic polynomial curves fitted to the individual coherence values in the sources identified by the ANOVA, suggest with both, three and six conditions, that the relationship between degradation levels and speech-tracking follows an inverted u-shape. These results nicely fit with fMRI findings of increased activation of (left) temporal and frontal inferior regions for degraded but yet intelligible stimuli compared to unaltered as well as completely unintelligible speech as reported by Davis and Johnsrude (2003) and interpreted as indicating recruitment of compensatory attentional resources. The authors showed that the effect in temporal areas was further depending on other acoustic features while the frontal regions did not respond to those suggesting that the frontal regions serve a more general executive function (Davis and Johnsrude 2003). Interestingly, those two regions (left inferior frontal gyrus and temporal region) exhibited enhanced fMRI responses to degraded but intelligible speech when attention was directed to the speech again interpreted as a marker of effortful listening (Wild et al. 2012) and left inferior cortex further plays a role in perceptual learning (Eisner et al. 2010). This is also consistent with a study showing non-native speakers produce higher delta/theta speech entrainment than native-speakers and the authors have also proposed this as reflecting the higher effort (Song and Iverson 2018). Similarly, speech tracking is increased during active compared to passive listening only for low levels of intelligibility (Vanthornhout et al. 2019). Further, the M50 of TRF is enhanced for degraded stimuli compared to unaltered ones in quiet environment as is delta entrainment, the latter again suggested to reflect listening efforts (Ding et al. 2014). Although, studies have also reported decreased theta entrainment for degraded speech (Ding et al. 2014; Rimmele et al. 2015; Peelle 2018), synchronization with the speech signal in both frequency bands is enhanced when attended to: multi-speaker and auditory spatial attention studies using sentences or narratives have repeatedly found stronger low-frequency (1-7 Hz) speech tracking for attended compared to unattended speech (Ding and Simon 2012; Horton et al. 2013; Zion Golumbic et al. 2013; Rimmele et al. 2015).

### 4.3 Alpha power decreases across degradation levels

Another commonly used measure in studies of degraded speech -a common operationalization for listening effort- is the alpha rhythm (Obleser and Weisz 2012; Obleser et al. 2012; McMahon et al. 2016; Miles et al. 2017). Interestingly, we found that alpha power followed a different pattern than coherence which became most obvious in study 2. While speech-tracking seems to have a u-shaped relationship with degradation level, alpha power shows a widespread decrease for the stimuli with less acoustic information compared to clear speech. Study 2 suggests that this decrease reaches a floor effect already with 5-channel vocoding. Both studies show the maximum of this decrease in left angular and parietal inferior gyrus. This is a region that has been reported to play a crucial role in complex speech comprehension (Van Ettinger-Veenstra et al. 2016), especially important in successful comprehension of degraded but predictable speech (Obleser et al. 2007; Hartwigsen et al. 2015) and importantly in perceptual learning of degraded speech (Eisner et al. 2010). The pattern of decreasing alpha power is further consistent with other studies using degradation of complex speech material as for example sentences (McMahon et al. 2016; Miles et al. 2017). However, studies using short and simple speech stimuli such as single words (Obleser and Weisz 2012; Becker et al. 2013) or digits (Obleser et al. 2012; Wöstmann et al. 2015) report enhanced alpha for stimuli with more acoustic detail compared to degraded sounds. This difference in findings might be linked to the linguistically more complex nature of the longer speech stimuli as also suggested by Miles and colleagues (2017).

To the best of our knowledge, so far no study investigated the influences of vocoded continuous speech on both alpha power and speech tracking. A study on a related topic found that cochlear implant (CI) users show alpha power to be positively correlated with subjective listening effort while speech-brain coherence showed a negative relationship (Dimitrijevic et al. 2019). Besides the differences in study groups (participants with normal hearing vs. CI users) and operationalisation of listening effort (vocoded speech vs. speech-in-noise-tasks) between our study and the one of Dimitrijevic et al. (2019), they used short auditory stimuli (digits) as many of the studies (Obleser and Weisz 2012; Obleser et al. 2012; Becker et al. 2013; Wöstmann et al. 2015) reporting the opposite pattern in alpha power than us.

### 4.4 Influences of stimulus material

Degrading the speech by vocoding as we did in the present study and as done by many other studies (e.g. Obleser et al. 2007; Miles et al. 2017) reduces the phonetic fine structure while temporal information, e.g. segmentation of syllables is preserved (Shannon et al. 1995). Based on our results, it seems that for challenging speech (reduced fine structure) people have to rely more on the temporal structure of speech leading to enhanced tracking in this frequency range (delta, theta) and that this process reverses beyond a critical point of degradation.

This effect might be amplified by the choice of long continuous speech stimuli. Unlike several other studies that used degraded single words (Obleser and Weisz 2012; Obleser et al. 2012; Becker et al. 2013; Wöstmann et al. 2015), we implemented long stimuli in the MEG studies, between 30 s and 3 min. The long duration of our stimuli might affect the perception of the different degradation levels differently via “warming-up” to the stimuli (Dorman et al. 1997). Experimental investigation of this warm-up or perceptual learning effect shows that indeed speech understanding increased over time for degraded stimuli (e.g. 4-chan vocoding: Rosen et al. 1999; 6-chan vocoding: Davis et al. 2005) and that this increase was bigger for sentences than for single words (Hervais-Adelman et al. 2008) and smallest for very strong (1-chan) or very little (24-chan) vocoding (Sohoglu and Davis 2016). Similar non-linear patterns have been reported for dual-task measures of listening effort. Reaction times (Wu et al. 2016) and pupil sizes (Zekveld and Kramer 2014) were enhanced for the middle range of speech intelligibility. Based on these findings, we speculate that the processes underlying listening to degraded speech dynamically vary depending on the stimulus length.

### 4.5 Can listening effort explain results?

Intuitively, listening effort seems like an easy-to-understand concept, and individuals usually can answer without difficulty whether listening to a stimulus was effortful. Stimulus degradation (e.g. vocoding) is a common operationalization for listening effort (e.g. Obleser and Weisz 2012). However, listening effort combines many dimensions. Peelle (2018) proposes a model comprising person-related characteristics (e.g. motivation) and stimulus-related characteristics (e.g. signal-to-noise ratio). Various measures exist for capturing listening effort, alpha power being one of them (e.g. Miles et al. 2017; Dimitrijevic et al. 2019). Speech tracking is not classically viewed as a measure of listening effort, nevertheless its modulations when listening to challenging speech have been interpreted as increased effort (Song and Iverson 2018) and increased attentional demands (Rimmele et al. 2015). Importantly, it has been shown that listening effort is of multidimensional nature with the different dimensions being captured by different measures that do not necessarily correlate (McGarrigle et al. 2014; Alhanbali et al. 2019). This fits with our findings of degradation levels (across a wide range) affecting alpha power and speech tracking differently and suggests that such measures are not ideal to explain abstract concepts as listening effort independent of circumstances.

### 4.6 Beyond listening effort, towards intelligibility

Results of the linear mixed models suggest that subjective intelligibility (behavioral response) can best be predicted by a combination of speech-coherence and alpha-power: We propose that for continuous degraded speech, understanding speech depends on the activation of veridical representations. Along the lines of a recent framework by Griffiths and colleagues (2019) we propose that this activation is reflected by alpha decrease. This process will however only support listening (e.g. reflected in the ability to track specific features) up to a specific (breaking) point, when speech becomes too degraded so that no veridical information is activated. This interpretation integrates well with the frameworks on alpha oscillations in the context of working memory as proposed for example by van Ede (2018) (2018), but also for auditory perception by Griffith et al. (2019), and for auditory memory by Kraft et al. (2019). Van Ede puts the idea forward that alpha power increases for tasks with sensory disengagement, while it decreases for tasks which recruit the sensory representation. Our task of asking participants to identify which of two presented words did occur within the just heard four last words of a speech stimulus will most likely recruit the sensory representation of words, thereby leading to a relative alpha decrease. For our results this would imply that the sensory representation is activated for all conditions of challenging speech as reflected by alpha decrease. For challenging conditions, this increased engagement is accompanied by increased tracking which decreases again when speech becomes unintelligible even though neural activation per se remains high. This fits nicely with The Ease of Language Understanding model (Rönnberg et al. 2013), which puts the idea forward, that the perceived phonological signals are tested against the stored phonological representation in memory and when they do not match, explicit working memory processes are elicited that aim at reconstructing the signal content. Several studies support the direct relationship between working memory and speech processing (e.g. Eisner et al. 2010; Rönnberg et al. 2010; Rudner et al. 2012). Within these frameworks, also different findings in the literature concerning alpha can be unified by taking the specific task and the resulting demands into account.

## 5. Conclusions

In sum, prior research reports mixed results concerning the link between degradation and speech-brain coherence as well as alpha power. We conducted two experiments with different levels of degradation, importantly of continuous speech. The results of these two studies show that the level of degradation affects speech tracking and alpha power differently: Speech tracking shows a u-shaped pattern with the easiest (original) and hardest (1-channel) to understand producing the lowest tracking values and the middle degradation level (5- channel) eliciting the highest tracking values. On the other hand, alpha power seems to overall decline with the declining clarity of speech. As study 2 shows, this decline likely reaches a floor effect also with 5-channel vocoding. Use of EEG signals is gaining momentum in the discussion about hearing aids improvement (Fiedler et al. 2016; Bech Christensen et al. 2018). In this context our findings have wider implications as they provide insights into more naturalistic, i.e. continuous, speech compared to single words and digits. Importantly, our results indicate that taking into account alpha modulations (interpreted in terms of neural activation) and neural speech tracking in a combined manner, may open up avenues to monitor the (subjective) intelligibility of speech sounds. This perspective goes beyond simplistic listening effort accounts and could have important applied implications.

## Acknowledgements

This study was supported by a FWF Einzelprojekt (P 31230). AK was supported by the Wellcome Trust [204820/Z/16/Z]. We would like to thank Joachim Gross and Hyojin Park for providing their original Matlab script for extracting the lip area. We would also like to thank Siri Ebert and Jonas Heilig for help with data acquisition.

## References

Alexandrou AM, Saarinen T, Kujala J, Salmelin R. 2018. Cortical entrainment: what we can learn from studying naturalistic speech perception. Lang Cogn Neurosci. 0:1–13.

Alhanbali S, Dawes P, Millman RE, Munro KJ. 2019. Measures of Listening Effort Are Multidimensional. Ear Hear. 40:1084–1097.

Bates D, Mächler M, Bolker B, Walker S. 2015. Fitting Linear Mixed-Effects Models Using lme4. J Stat Softw. 67:1–48.

Bech Christensen C, Hietkamp RK, Harte JM, Lunner T, Kidmose P. 2018. Toward EEG-Assisted Hearing Aids: Objective Threshold Estimation Based on Ear-EEG in Subjects With Sensorineural Hearing Loss. Trends Hear. 22.

Becker R, Pefkou M, Michel CM, Hervais-Adelman AG. 2013. Left temporal alpha-band activity reflects single word intelligibility. Front Syst Neurosci. 7.

Benjamini Y, Hochberg Y. 1995. Controlling the False Discovery Rate: A Practical and Powerful Approach to Multiple Testing. J R Stat Soc Ser B Methodol. 57:289–300.

Brainard DH. 1997. The Psychophysics Toolbox. Spat Vis. 10:433–436.

Chandrasekaran C, Trubanova A, Stillittano S, Caplier A, Ghazanfar AA. 2009. The natural statistics of audiovisual speech. PLoS Comput Biol. 5.

Crosse MJ, Di Liberto GM, Bednar A, Lalor EC. 2016. The Multivariate Temporal Response Function (mTRF) Toolbox: A MATLAB Toolbox for Relating Neural Signals to Continuous Stimuli. Front Hum Neurosci. 10.

Davis MH, Johnsrude IS. 2003. Hierarchical processing in spoken language comprehension. J Neurosci Off J Soc Neurosci. 23:3423–3431.

Davis MH, Johnsrude IS, Hervais-Adelman A, Taylor K, McGettigan C. 2005. Lexical Information Drives Perceptual Learning of Distorted Speech: Evidence From the Comprehension of Noise-Vocoded Sentences. J Exp Psychol Gen. 134:222–241.

Dimitrijevic A, Smith ML, Kadis DS, Moore DR. 2019. Neural indices of listening effort in noisy environments. Sci Rep. 9:1–10.

Ding N, Chatterjee M, Simon JZ. 2014. Robust Cortical Entrainment to the Speech Envelope Relies on the Spectro-temporal Fine Structure. NeuroImage. 88:41–46.

Ding N, Simon JZ. 2011. Neural coding of continuous speech in auditory cortex during monaural and dichotic listening. J Neurophysiol. 107:78–89.

Ding N, Simon JZ. 2012. Emergence of neural encoding of auditory objects while listening to competing speakers. Proc Natl Acad Sci. 109:11854–11859.

Dorman MF, Loizou PC, Rainey D. 1997. Speech intelligibility as a function of the number of channels of stimulation for signal processors using sine-wave and noise-band outputs. J Acoust Soc Am. 102:2403–2411.

Ede F van. 2018. Mnemonic and attentional roles for states of attenuated alpha oscillations in perceptual working memory: a review. Eur J Neurosci. 48:2509–2515.

Eisner F, McGettigan C, Faulkner A, Rosen S, Scott SK. 2010. Inferior Frontal Gyrus Activation Predicts Individual Differences in Perceptual Learning of Cochlear-Implant Simulations. J Neurosci. 30:7179–7186.

Fiedler L, Obleser J, Lunner T, Graversen C. 2016. Ear-EEG allows extraction of neural responses in challenging listening scenarios — A future technology for hearing aids? Presented at the Conference proceedings: … Annual International Conference of the IEEE Engineering in Medicine and Biology Society. IEEE Engineering in Medicine and Biology Society. Conference. p. 5697–5700.

Frey JN, Mainy N, Lachaux J-P, Muller N, Bertrand O, Weisz N. 2014. Selective Modulation of Auditory Cortical Alpha Activity in an Audiovisual Spatial Attention Task. J Neurosci. 34:6634–6639.

Gaudrain, E. 2016. Vocoder, v1.0.

Greenberg S. 1998. A syllable-centric framework for the evolution of spoken language. Behav Brain Sci. 518.

Griffiths BJ, Mayhew SD, Mullinger KJ, Jorge J, Charest I, Wimber M, Hanslmayr S. 2019. Alpha/beta power decreases track the fidelity of stimulus-specific information. eLife. 8:e49562.

Gross J, Hoogenboom N, Thut G, Schyns P, Panzeri S, Belin P, Garrod S. 2013. Speech Rhythms and Multiplexed Oscillatory Sensory Coding in the Human Brain. PLoS Biol. 11.

Hartwigsen G, Golombek T, Obleser J. 2015. Repetitive transcranial magnetic stimulation over left angular gyrus modulates the predictability gain in degraded speech comprehension. Cortex, Special issue: Prediction in speech and language processing. 68:100–110.

Hauswald A, Lithari C, Collignon O, Leonardelli E, Weisz N. 2018. A Visual Cortical Network for Deriving Phonological Information from Intelligible Lip Movements. Curr Biol CB. 28:1453–1459.e3.

Hervais-Adelman A, Davis MH, Johnsrude IS, Carlyon RP. 2008. Perceptual learning of noise vocoded words: Effects of feedback and lexicality. J Exp Psychol Hum Percept Perform. 34:460–474.

Horton C, D’Zmura M, Srinivasan R. 2013. Suppression of competing speech through entrainment of cortical oscillations. J Neurophysiol. 109:3082–3093.

Howard MF, Poeppel D. 2010. Discrimination of Speech Stimuli Based on Neuronal Response Phase Patterns Depends on Acoustics But Not Comprehension. J Neurophysiol. 104:2500–2511.

Keitel A, Gross J, Kayser C. 2018. Perceptually relevant speech tracking in auditory and motor cortex reflects distinct linguistic features. PLoS Biol. 16.

Kleiner M, Brainard D, Pelli D, Ingling A, Murray R, Broussard C. 2007. What’s new in psychtoolbox-3. Perception. 36:1–16.

Kraft N, Demarchi G, Weisz N. 2019. Auditory cortical alpha desynchronization prioritizes the representation of memory items during a retention period. bioRxiv. 626929.

Loizou PC, Dorman M, Tu Z. 1999. On the number of channels needed to understand speech. J Acoust Soc Am. 106:2097–2103.

Luo H, Poeppel D. 2007. Phase Patterns of Neuronal Responses Reliably Discriminate Speech in Human Auditory Cortex. Neuron. 54:1001–1010.

Maris E, Oostenveld R. 2007. Nonparametric statistical testing of EEG- and MEG-data. J Neurosci Methods. 164:177–190.

Mattout J, Henson RN, Friston KJ. 2007. Canonical source reconstruction for MEG. Comput Intell Neurosci. 2007.

McGarrigle R, Munro KJ, Dawes P, Stewart AJ, Moore DR, Barry JG, Amitay S. 2014. Listening effort and fatigue: what exactly are we measuring? A British Society of Audiology Cognition in Hearing Special Interest Group “white paper.” Int J Audiol. 53:433–440.

McGettigan C, Faulkner A, Altarelli I, Obleser J, Baverstock H, Scott SK. 2012. Speech comprehension aided by multiple modalities: behavioural and neural interactions. Neuropsychologia. 50:762–776.

McMahon CM, Boisvert I, de Lissa P, Granger L, Ibrahim R, Lo CY, Miles K, Graham PL. 2016. Monitoring Alpha Oscillations and Pupil Dilation across a Performance-Intensity Function. Front Psychol. 7.

Miles K, McMahon C, Boisvert I, Ibrahim R, de Lissa P, Graham P, Lyxell B. 2017. Objective Assessment of Listening Effort: Coregistration of Pupillometry and EEG. Trends Hear. 21:2331216517706396.

Nolte G. 2003. The magnetic lead field theorem in the quasi-static approximation and its use for magnetoencephalography forward calculation in realistic volume conductors. Phys Med Biol. 48:3637–3652.

Obleser J, Weisz N. 2012. Suppressed alpha oscillations predict intelligibility of speech and its acoustic details. Cereb Cortex. 22:2466–2477.

Obleser J, Wise RJS, Dresner MA, Scott SK. 2007. Functional Integration across Brain Regions Improves Speech Perception under Adverse Listening Conditions. J Neurosci. 27:2283–2289.

Obleser J, Wöstmann M, Hellbernd N, Wilsch A, Maess B. 2012. Adverse Listening Conditions and Memory Load Drive a Common Alpha Oscillatory Network. J Neurosci. 32:12376–12383.

Oostenveld R, Fries P, Maris E, Schoffelen JM. 2011. FieldTrip: Open source software for advanced analysis of MEG, EEG, and invasive electrophysiological data. Comput Intell Neurosci. 2011.

Peelle JE. 2018. Listening Effort: How the Cognitive Consequences of Acoustic Challenge Are Reflected in Brain and Behavior. Ear Hear. 39:204.

Pelli DG. 1997. The VideoToolbox software for visual psychophysics: transforming numbers into movies. Spat Vis. 10:437–442.

Poeppel D. 2003. The analysis of speech in different temporal integration windows: cerebral lateralization as ‘asymmetric sampling in time.’ Speech Commun, The Nature of Speech Perception. 41:245–255.

Riecke L, Formisano E, Sorger B, Başkent D, Gaudrain E. 2018. Neural Entrainment to Speech Modulates Speech Intelligibility. Curr Biol. 28:161–169.e5.

Rimmele JM, Zion Golumbic E, Schröger E, Poeppel D. 2015. The effects of selective attention and speech acoustics on neural speech-tracking in a multi-talker scene. Cortex, Special issue: Prediction in speech and language processing. 68:144–154.

Rönnberg J, Lunner T, Zekveld A, Sörqvist P, Danielsson H, Lyxell B, Dahlström Ö, Signoret C, Stenfelt S, Pichora-Fuller MK, Rudner M. 2013. The Ease of Language Understanding (ELU) model: theoretical, empirical, and clinical advances. Front Syst Neurosci. 7.

Rönnberg J, Rudner M, Lunner T, Zekveld AA. 2010. When cognition kicks in: working memory and speech understanding in noise. Noise Health. 12:263–269.

Rosen S, Faulkner A, Wilkinson L. 1999. Adaptation by normal listeners to upward spectral shifts of speech: Implications for cochlear implants. J Acoust Soc Am. 106:3629–3636.

Rudner M, Lunner T, Behrens T, Thorén E, Rönnberg J. 2012. Working Memory Capacity May Influence Perceived Effort during Aided Speech Recognition in Noise. J Am Acad Audiol. 23:577–589.

Shannon RV, Zeng F-G, Kamath V, Wygonski J, Ekelid M. 1995. Speech Recognition with Primarily Temporal Cues. Science. 270:303–304.

Smith ZM, Delgutte B, Oxenham AJ. 2002. Chimaeric sounds reveal dichotomies in auditory perception. Nature. 416:87–90.

Sohoglu E, Davis MH. 2016. Perceptual learning of degraded speech by minimizing prediction error. Proc Natl Acad Sci. 113:E1747–E1756.

Song J, Iverson P. 2018. Listening effort during speech perception enhances auditory and lexical processing for non-native listeners and accents. Cognition. 179:163–170.

Strelnikov K, Massida Z, Rouger J, Belin P, Barone P. 2011. Effects of vocoding and intelligibility on the cerebral response to speech. BMC Neurosci. 12.

Van Ettinger-Veenstra H, McAllister A, Lundberg P, Karlsson T, Engström M. 2016. Higher Language Ability is Related to Angular Gyrus Activation Increase During Semantic Processing, Independent of Sentence Incongruency. Front Hum Neurosci. 10.

Van Veen BD, van Drongelen W, Yuchtman M, Suzuki A. 1997. Localization of brain electrical activity via linearly constrained minimum variance spatial filtering. IEEE Trans Biomed Eng. 44:867–880.

Vanthornhout J, Decruy L, Francart T. 2019. Effect of Task and Attention on Neural Tracking of Speech. Front Neurosci. 13:977.

Viswanathan V, Bharadwaj HM, Shinn-Cunningham BG. 2019. Electroencephalographic Signatures of the Neural Representation of Speech during Selective Attention. eneuro. 6:ENEURO.0057-19.2019.

Wild CJ, Yusuf A, Wilson DE, Peelle JE, Davis MH, Johnsrude IS. 2012. Effortful Listening: The Processing of Degraded Speech Depends Critically on Attention. J Neurosci. 32:14010–14021.

Wöstmann M, Herrmann B, Wilsch A, Obleser J. 2015. Neural Alpha Dynamics in Younger and Older Listeners Reflect Acoustic Challenges and Predictive Benefits. J Neurosci. 35:1458–1467.

Wu Y-H, Stangl E, Zhang X, Perkins J, Eilers E. 2016. Psychometric functions of dual-task paradigms for measuring listening effort. Ear Hear. 37:660–670.

Zekveld AA, Kramer SE. 2014. Cognitive processing load across a wide range of listening conditions: Insights from pupillometry. Psychophysiology. 51:277–284.

Zion Golumbic EM, Ding N, Bickel S, Lakatos P, Schevon CA, McKhann GM, Goodman RR, Emerson R, Mehta AD, Simon JZ, Poeppel D, Schroeder CE. 2013. Mechanisms Underlying Selective Neuronal Tracking of Attended Speech at a “Cocktail Party.” Neuron. 77:980–991.

Zoefel B, Archer-Boyd A, Davis MH. 2018. Phase Entrainment of Brain Oscillations Causally Modulates Neural Responses to Intelligible Speech. Curr Biol. 28:401–408.e5.

